# Continuous State HMMs for Modeling Time Series Single Cell RNA-Seq Data

**DOI:** 10.1101/380568

**Authors:** Chieh Lin, Ziv Bar-Joseph

## Abstract

**Motivation:** Methods for reconstructing developmental trajectories from time series single cell RNA-Seq (scRNA-Seq) data can be largely divided into two categories. The first, often referred to as pseudotime ordering methods, are deterministic and rely on dimensionality reduction followed by an ordering step. The second learns a probabilistic branching model to represent the developmental process. While both types have been successful, each suffers from shortcomings that can impact their accuracy.

**Results:** We developed a new method based on continuous state HMMs (CSHMMs) for representing and modeling time series scRNA-Seq data. We define the CSHMM model and provide efficient learning and inference algorithms which allow the method to determine both the structure of the branching process and the assignment of cells to these branches. Analyzing several developmental single cell datasets we show that the CSHMM method accurately infers branching topology and correctly and continuously assign cells to paths, improving upon prior methods proposed for this task. Analysis of genes based on the continuous cell assignment identifies known and novel markers for different cell types.

**Availability:** Software and Supporting website: www.andrew.cmu.edu/user/chiehll/CSHMM/

**Supplementary information:** Supplementary data are available at *Bioinformatics* online.

## 1 Introduction

The ability to profile gene expression and other genomic data in single cells has already led to several new findings. Using single cell expression data (scRNA-Seq) researchers can better identify cell specific pathways and genes which are often missed when profiling cell mixtures. scRNA-Seq analysis of developmental programs, various tissues and perturbations has already identified new cell types, new pathways and new marker genes for a variety of biological systems and conditions (Trapnell et al., 2014; Shalek et al., 2013; Treutlein et al., 2014).

Several scRNA-Seq studies have profiled time series data, most notably during development of various organs and systems (Treutlein et al., 2014, 2016). A key question that emerges in such studies is the ability to connect different cell types over time. Unlike experiments that profile bulk samples (or population of cells), in which a sample at time point *t* + 1 is assumed to arise from the sample at time *t* (Bar-Joseph et al., 2012), in single cell studies it is not always clear what cell type in time *t* led to a cell being profiled in time *t* + 1. Since scRNA-Seq studies fully consume the cell (which effectively makes it a snapshot), it is not possible to trace it over time which make it difficult to connect progenitor cells to their descendents, or to follow the response of specific cell types over time. In addition, in many single cell studies cells are not completely synchronized within a sample and so cells measured at a specific time point may be more similar to cells at other time point (in terms of their perceived developmental or differentiation time) which may require the reassignment of cells between the measured time points.

To address these issues, a number of methods, often referred to as pseudotime inference methods, have been proposed (Trapnell et al., 2014; Qiu et al., 2017; Setty et al., 2016; Rizvi et al., 2017). These methods order cells along a transcriptomic trajectory in embedded space such that cells that are close in that space are also assumed to be close in terms of their biological states. By tracing paths and trajectories of pseudotemporally ordered cells these methods determine the set of states leading from the starting point to the (often differentiated) final cell fate. Pseudotime and other tools developed for the analysis of time series scRNA-Seq data can be largely divided based on the method they used (probabilistic or deterministic) and the representation they provide (continuous vs. discrete cell assignments). Deterministic methods utilize dimensionality reduction (often to two components) to obtain a graph representation of all cells in a lower dimension embedding and then rely on graph analysis (usually an extension or variant of minimum spanning trees) or other modeling methods (for example, Gaussian Processes (GP)) to connect cells in order to obtain a continuous trajectory for the cells in the study. Some methods that use variants of this strategy include DPT (Haghverdi et al., 2016), scTDA (Rizvi et al., 2017), PCA analysis (Treutlein et al., 2014), Monocle 2 (Trapnell et al., 2014), Wanderlust (Bendall et al., 2014)) and GPLVM (Reid and Wernisch, 2016; Lönnberg et al., 2017). In contrast, probabilistic state methods assign cells to a discrete (and often small) number of states in a probabilistic graphical model and determine trajectories based on the graph structure. Methods that use this strategy include SCUBA(Marco et al., 2014) and TASIC (Rashid et al., 2017). While both types of methods have been successful in analyzing various types of time series scRNA-Seq data, they both suffer from problems that limit their general use. Deterministic methods often do not take into account noise, which is very prevalent in scRNA-Seq data (Buettner et al., 2015; Shapiro et al., 2013). In addition, by reducing the dimensions to a very small number of components they effectively ignore much of the profiled data. Finally, some of these methods cannot infer more than two branches in the trajectory which is often not enough for developmental and other studies. Probabilistic methods overcome these issues by introducing transitions that handle branching and emissions that can handle noise. However, the resulting graph from these methods includes only a small number of states which makes it hard to infer continuous trajectories of genes along the developmental process and also forces cells that can be pretty distant in the time they represent to the same state.

Here we present a new method for ordering cells in scRNA-Seq studies which combines the continuous representation offered by the deterministic methods and the ability to handle noise provided by the probabilistic methods. Our algorithm is based on the use of Continuous State HMMs (CSHMMs) (Ainsleigh, 2001). Unlike standard HMMs which are defined using a discrete set of states, continuous state HMMs can have infinitely many states and so cells can be assigned to a much more detailed trajectory. We discuss how to formulate the CSHMMs for scRNA-Seq data and how to perform learning and inference in this model. Once we learn a CSHMM model all cells are assigned to specific locations along paths which allows users to associate cells with specific fates and to reconstruct continuous developmental trajectories for the genes along each path. We applied our CSHMM to several scRNA-Seq datasets. As we show, the method was able to correctly assign cells to paths in order to reconstruct developmental trajectories for these processes improving upon the models obtained by both the deterministic and prior probabilistic models. Using the learned cell assignment we were also able to identify several novel genes for the different cell fate trajectories.

## 2 Methods

### 2.1 Dataset processing

We collected time-series mouse scRNA-Seq data for lung (Treutlein et al., 2014) and for mouse embryonic fibroblasts (MEFs) re-programming to neurons (Treutlein et al., 2016). Both datasets were processed in a similar way to the processing performed in the original paper: We removed genes with FPKM < 1 in all cells and genes with zero variance. Next expression values were transformed to log FPKM. We also analyzed a zebrafish embryogenesis scRNA-Seq dataset (Farrell et al., 2018). This dataset is log TPM and genes expressed in less then 5% of cells are removed. The processed mouse lung data consisted of 152 cells with 15K genes and 3 time points (E14.5, E16.5, E18.5), the mouse MEF reprogramming data consisted of 252 cells with 12K genes and 4 time points (0, 2, 5, 22 days), the zebrafish data consisted of 38731 cells with 6K genes and 12 time points (from 3.3 to 12 hours).

### 2.2 CSHMM model formulation

Fig. 1 presents the CSHMM model structure. HMMs define a transition probability between states and emission probability for each state. CSHMMs defines the same set of parameters. However, since they have infinite many states (in our case corresponding to continuous time) both transition and emission probabilities are a function of the specific *path* a state resides on. Split points represent time points where we allow cells to split into different lineages and paths are defined as the collection of (infinitely many) states between two such split events. Note that in our model we learn the location of the splits from data and while these are initialized with the sampling rate (i.e. initially we use the sampled time points to define the split locations) as we discuss below the model can add splits between two time points to account for the asynchronous nature of cells in some studies.

**Fig. 1.**
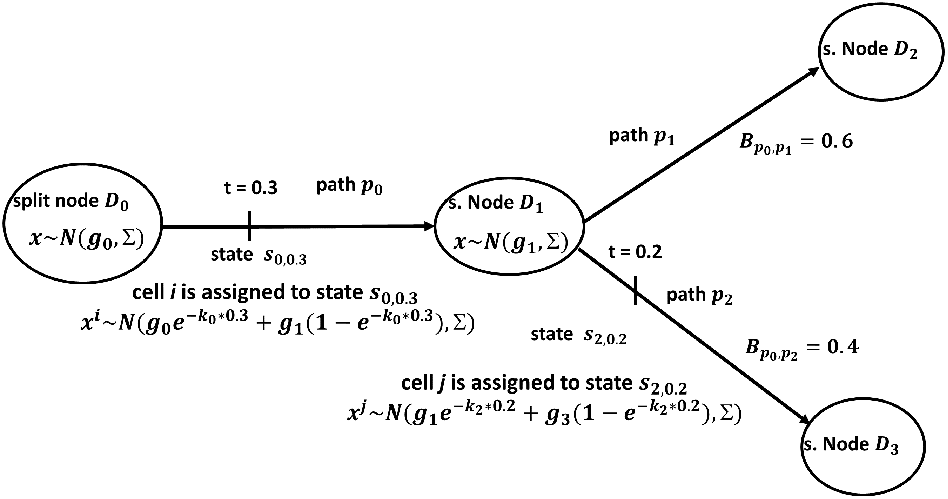
CSHMM model structure and parameters. Each path represents a set of infinite states parameterized by the path number and the location along the path. For each such state we define an emission probability and a transition probability to all other states in the model. Emission probability for a gene along a path is a function of the location of the state and a gene specific parameter *k* which controls the rate of change of its expression along the path. Split nodes are locations where paths split and are associated with a branch probability. Each cell is assigned to a state in the model. See text for complete details

Each cell is assigned to a specific state along one of the paths which corresponds to both, the time inferred for it by the algorithm and the cell type it belongs to. In addition to the state assignment and transitions at split nodes the model also encodes emission probabilities. Following prior work on modeling expression with HMMs (Lin et al., 2008) we use a Gaussian emission model and assume independence for gene specific expression levels conditioned on the state. To define an emission probability for a state we use the *relative* location of a state along a specific path. We define a state by the path number and the relative time for this path. We denote by *s_p,t_* the state representing time 0 ≤ *t* ≤ 1 on path *p*(*D_a_* → *D_b_*), where *a, b* are the indices of the split nodes. Let *i* be a cell assigned to *s_p,t_*. We denote by 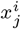 the expression of gene *j*. The emission probability for gene *j* in cell *i* assigned to state *s_p,t_* is thus assumed to be 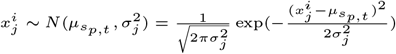 Where

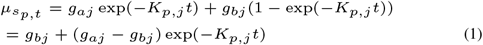

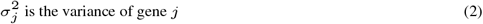

Here, *g_a_j* is the mean expression for gene j at split node a. We assume a continuous change in expression for a subset of the genes along a path. However, we want to learn the specific shape of the change curve and so we add a parameter *K_p,j_* which controls the rate of change for gene j on path p, allowing different genes to change at different rates. Using these notations we next define the following parameters that are required to specify a CSHMM: λ = (*V, π, S, A, E*), where all the symbol definitions are presented in Table 1.

**Table 1.**
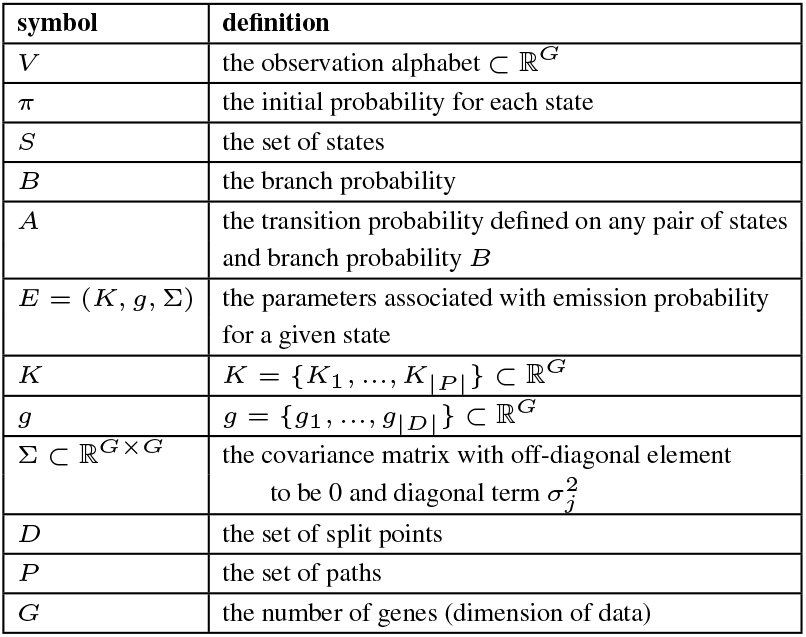
The parameter definition for CSHMM

Each cell *i* is associated with an expression vector *X^i^* ∈ ℝ^*G*^, and a (hidden) state *y^i^* = *s_p,t_*. The observation alphabet *V* ⊂ ℝ^*G*^, is thus a real value vector with dimension |*G*|, where G are the set of genes in our input set. We associate a root state so,o with each HMM with initial probability of 1 (*π_p,t_* = 1 for state *s*_0,0_ and *π_p,t_* =0 for all other states). The transition probability *A*(*s*_*p*1, *t*_1__, *s*_*p*2, *t*_2__) for each pair of states *s*_*p*1, *t*_1__, *s*_*p*2, *t*_2__ ∈ *S* is defined as follows:

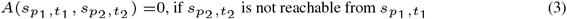

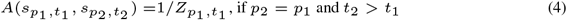

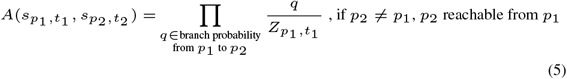

Where *Z*_*p*1, *t*_1__ is a normalizing factor for the transition probability going out of state *s*_*p*1, *t*_1__ i.e..

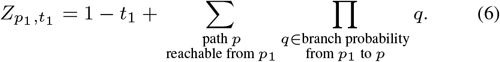

The branch probability is defined on split nodes as shown in Fig. 1. The use of branching probabilities leads to lower likelihood for cell assignments to later (more specific) paths in the branching tree. This is similar to prior probabilistic methods for reconstructing branching trajectories (Rashid et al., 2017). The idea here is that earlier stages are often less specific (higher entropy (Teschendorff and Enver, 2017), while later stages (representing specific fates) have a tighter expression profile. Thus, cells that represent specific cell types will still be assigned to their correct (late) stage based on their expression profile while noisier cells would be assigned to the earlier stages.

To see that this is indeed a Continuous-State Hidden Markov Model (CSHMM) model we note that the model contains a continuous set of states with well defined emission and transitions probabilities (transition probabilities integrate to 1 for each state). Transitions and emissions only depend on the current state. Each observation is assumed to have been emitted from one of the states in the model.

Since we cannot assume that the time stamp associated with each cell is the correct time (to account for asynchrony) we need to determine cell assignments. In addition, we do not know the structure of the model in advance. We thus developed an Expectation Maximization (EM) algorithm which can jointly infer the model structure, parameters and cell assignments.

### 2.3 Likelihood function for the CSHMM model

Since CSHMMs are probabilistic models, to determine the optimal structure and parameters we first need to define the likelihood function that the model is trying to optimize. Denote by *X^i^* the expression profile of cell *i*. Let 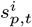 denote the (unobserved) state which ‘emitted’ the expression of cell *i* (i.e. the state to which cell *i* is assigned to). Given an expression input matrix *X* = {*X*^1^,…, *X^N^*} and hidden variables *Y* = {*y*^1^,…, *y^N^*)} where 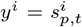 is the state for cell i we can write the log likelihood as follows:

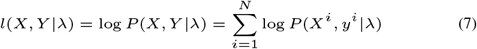

Which can be can further decomposed using the parameters described above as:

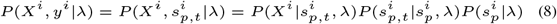

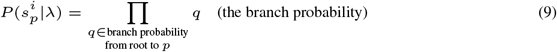

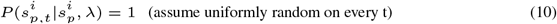

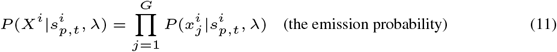

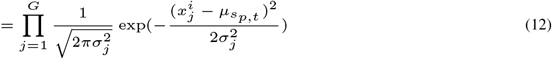

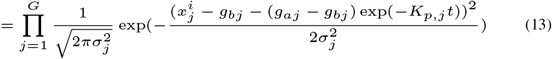

Thus, the complete log likelihood for *N* input cells is:

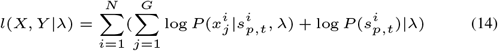

Note that in equation 10, 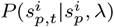 is a probability density function over path *p* with domain 0 ≤ *t* ≤ 1. The normalization performed in equations 3 – 5 guarantees that all transition probabilities for a state integrates to 1.

#### 2.3.1 Constraining expression changes along a path

Similar to prior pseudotime ordering methods our algorithm relies on the assumption that cells that are close to each other along the developmental trajectory have a similar (though not identical) expression profile. This implies that for most genes we would expect to see relatively small changes in expression whereas for a few genes (which may define the changes that the cell undergoes during the process) we expect larger changes. Thus, we expect differences between the expression profiles of consecutive split nodes to be sparse. To encode our assumption about the sparseness of the difference vector Δ*g* we use *L*_1_ regularization on the difference. Minimizing the *L*_1_ regularization term for negative log-likelihood (NLL) is equivalent to maximizing the complete likelihood multiplied by the Laplace prior distribution (Tibshirani, 1996). The Laplace prior distribution on Δ*g* and parameter *h* is:

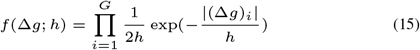

where *h* > 0 is the scale of the distribution.

Adding this regularization, the log likelihood function changes to:

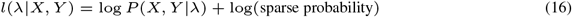

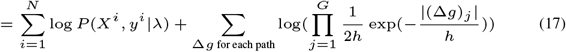

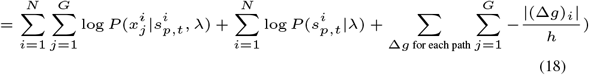

### 2.4 Model initialization

For model initialization we slightly modify the strategy used in (Ding et al., 2018). We construct an initial cell differentiation tree by clustering the cells, and then compute the distance of each of the clusters to the root of the tree (cells in first time point). Using this distance function clusters are assigned to different levels in the tree (where clusters in each level are significantly more distant from the root than the preceding level). Finally, we connect each cluster (except the root cluster) at level i to a parent cluster in level *i* – 1 by selecting the closest cluster, in expression space, in level *i* – 1. See Supporting methods for complete details. Following this initialization step each cluster is associated with a path (the edge connecting it to its parent). Finally, cells in each cluster are randomly assigned along the path for that cluster. Split nodes are defined for cases where two or more clusters at a specific level connect to the same cluster at the level above them.

### 2.5 Learning and Inference (EM algorithm)

We use an EM algorithm to learn the parameters of the model and to infer new cell assignment. Given initial cell assignments, the branching probabilities can be easily inferred using standard Maximum Likelihood Estimation (supporting methods). In the supplement we discuss how to learn the emission probability parameters which, due to the *K* parameter requires an optimization of a non convex target function. As for cell assignment, given model parameters we assign each cell to a state *s_p,t_* which maximizes the log-likelihood of the resulting model. Again, since the likelihood function is not concave, determining a optimal value t for a cell assigned to path p is challenging. In the supplement we discussed a sampling strategy for solving this problem which we use to assign cells.

### 2.6 Modifying the model structure

So far we assumed a fixed model structure. However, as part of the EM algorithm cells are re-assigned and so some paths that started with several cells may become empty while for others we may need to reassign their parents as their expression parameters change. To allow for structure changes during the learning process we do the following. Following each EM iteration, we test for two things: First, if a path has less than 3 cells assigned to it we remove it from the model and connect any following paths to the path parents. In addition, we allow the algorithm to connect split nodes to different parents in the level above them. For this, we try to connect every path at a certain level to all paths at the prior level it was not connected to. For each such new connection we re-compute the log-likelihood for all cells assigned on the path. If the log likelihood increases for this set of cells we keep the new relationship, otherwise we do not. This is repeated for every possible connection resulting in the structure that maximizes the likelihood for the current assignments we have.

### 2.7 Analysis of gene expression for specific cell fates

To determine the set of genes associated which specific fates (a set of paths from root to a leaf in the model), we calculate the Spearman correlation between their expression values and the ordering of the cells assigned to the set of paths leading to a specific fate. We use gprofiler (Reimand et al., 2016) for GO of the top 300 genes. For plotting gene expression we use a 4 degree polynomial to interpolate expressions in the different cells assigned to a trajectory. For each leaf node, we scale all cell assignments between the root and the node to be between 0 and 1 so that all expression profiles are plotted with the same length.

## 3 Results

To test the CSHMM model and to compare the results to prior pseudotime ordering methods we used several time series scRNA-Seq datasets. The first dataset is for mouse lung development (Treutlein et al., 2014). After preprocessing (Methods), the lung dataset consisted of 152 cells with 15K genes, measured at 3 time points (14.5, 16.5, 18.5 days). Cells at time point 18.5 are labeled with one of the following cell types: alveolar type 1 (AT1), alveolar type 2 (AT2), bipotential progenitor (BP), Clara and Ciliated. Cells at earlier time points were not labeled in the original paper. We label these cells as NA_14 or NA_16 based on their time point. The second dataset profiled the process in which mouse embryonic fibroblasts (MEFs) are induced to become neuronal (iN) cells (Treutlein et al., 2016)

This data contained 4 time points (0, 2, 5, 22 days) starting with MEF cells at day 0. Using known markers, Day 22 cells were labeled in the original paper with one of the following cell types: Neuron, Myocyte, Fibroblast. For the rest of the cells we used the assignments in the original papers for the plots, though they were not used by the CSHMM algorithm. In addition to these two well annotated, but rather small, datasets we also tested the CSHMM on a much larger zebrafish embryogenesis dataset (Farrell et al., 2018). This dataset has close to 40,000 cells profiled at 12 time points (from 3.3 to 12 hours). Cells in the last time point (only) were labeled with one of 25 cell types based on marker genes.

### 3.1 Application of CSHMM to lung developmental data

Fig. 2 presents the resulting CSHMM branching model for the lung development data and the distribution of cells along its paths (based on the state assigned by the model). As can be seen, the CSHMM method was able to assign different cell types to different paths correctly, for example, ciliated (path 2), Clara (path 3), AT1 (path 7), and AT2 (path 5) are all correctly associated with a terminal path. The bi-progenitor (BP) cells (path 4) are mostly assigned to the predecessors of the AT1 and AT2 paths in agreement with prior observations (Treutlein et al., 2014). This highlights the ability of the method to assign cells measured at the same time point (E18.5) to different times in the model. The ability to correctly reconstruct the branching trajectory for such *in vivo* data is not trivial. As we show in Fig. 3 (a)(b)(c)(d), dimensionality reduction based methods that have been used in the past for pseudo time ordering, including PCA (Treutlein et al., 2014), TSNE (for which we used the optimal parameters, Supporting Methods), GPLVM following PCA (Campbell and Yau, 2016), and Monocle 2 ^1^ (Trapnell et al., 2014; Qiu et al., 2017) where unable to fully reconstruct the known developmental trajectory for this data.

**Fig. 2.**
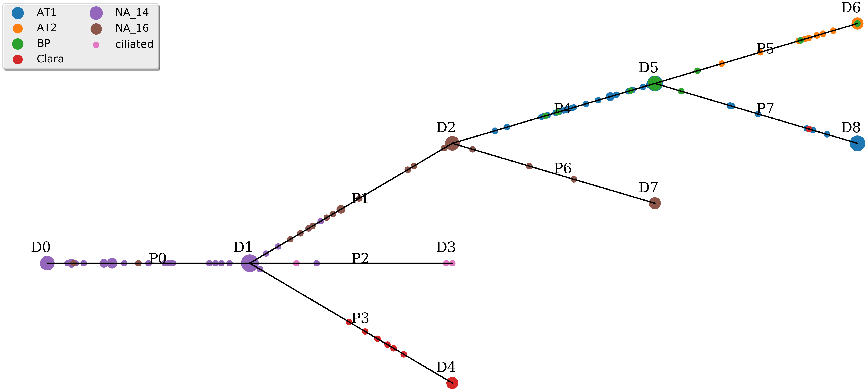
CSHMM model structure and continuous cell assignment for the lung developmental dataset. D nodes are split nodes and P edges are paths as shown in Fig. 1. Each small circle is a cell assigned to a state on the a path. The bigger the circle the more cells are assigned to this state. Cells are colored based on the cell type / time point assigned to them in the original paper.

PCAis able to identify clusters for different cell types but the projection of the reduced dimensional cells cannot reconstruct the known trajectory over time. Similarly, TSNE was also unable to separate some cell types for the later time point and was mixing E14.5 and E16.5 cells. GPLVM correctly orders E14.5 and E16.5 cells, however, it is unable to determine branching models for the different cells types in E18.5 and is also unable to determine the relative earlier ordering of the BP cells. Monocle 2 was able to generate trajectories, associating cells with specific time points, however, for this data it finds only 1 split point and was also unable to correctly separate the E18.5 cells according to their types. We also tried to compare to scTDA (Rizvi et al., 2017), however that method requires a commercial software from Ayasdi Inc. that we did not have access to. We have also compared the results to prior probabilistic methods that use a discrete set of states (Rashid et al., 2017). For this we have re-run the CSHMM algorithm but this time allowing cells to be assigned only to the endpoints of paths themselves and not to intermediate points. Results are presented in Supporting Fig. 1. As can be seen, although the discrete version leads to good result in terms of cell assignments, there are some differences. Specifically, BP cells are mostly assigned to terminal paths in these models, rather than intermediate paths. Further, as we show below, the CSHMM model is better at identifying cell type specific genes when compared to prior probabilistic discrete models.

#### 3.1.1 Identifying cell type specific genes in the lung dataset

The continuous nature of the CSHMM allows us to reconstruct the full gene expression trajectories for each path / cell type (ending at a leaf in our model). For this we use the ordering of cells from root to leaf for each of the leaves. To overcome noise in individual cell measurements we fit a continuous function (a polynomial of degree 4) to the set of values for each gene and plot the resulting curve. We then use these curves to search for genes that are specifically correlated with a leaf (cell type, Methods). In addition, we can compare trajectories for genes between two leafs to identify genes that are uniquely associated with one cell type. To illustrate the advantages of such analysis we have plotted in Fig. 4 (a) the trajectories of some of the known markers for the cell types in the data and additional genes (Fig. 4 (b)) that, while not currently known as markers or cell type specific, are expressed in a similar manner to known markers and so are predicted to be novel markers for specific cell types. For example, Aqp5’s expression is high for the AT1 path, but is strongly decreasing in the AT2 path after the split between these two paths. For the novel markers, Bc051019 displays very similar expression to the known cillated marker Foxj1. An independent study profiling scRNA-Seq of lung epithelial cells (Du et al., 2015) has also identified it as a ciliated marker. See Supporting Table 1 for a full list of genes that are significantly associated with each pair of paths. The ability to reconstruct the full trajectory of the genes along the paths based on the continuous state assignments is also an advantage of the CSHMM compared to prior methods. As we show in Fig. 4 (c), for several genes the trajectory assigned by the CSHMM model is more accurate (based on known biology) than their trajectory in a discrete HMM model. For example, Etv5 and Soat1 are two AT1/AT2 markers found by CSHMM which are not identified by a regular HMM. Recent studies suggest that Etv5 is essential for the maintenance of AT2 cells (Zhang et al., 2017), and Soat1 is expressed in AT2 cells (Gutierrez et al., 2016). We have also performed GO analysis on the set of genes that are identified for each leaf path (See supporting website). Several of the functions identified agree with known functions for the terminal paths. For example, the most significant GO category for genes correlated with path 2 the *cillated path* was cilium assembly (p-value = 1e-14) which is indeed the major function of cillated cells. Similarly, epithelium development was one of the top categories for path 3 *Clara path* (p-value = 4e-6). For path 7 (AT1 cells) the top categories were related to extracellular matrix (p-value = 3e-9), which is known to be associated with the development of this cell type (Olsen et al., 2005).

### 3.2 Application of CSHMM to neural developmental data

We have also analyzed a slightly more complicated MEF cell differentiation dataset (Treutlein et al., 2016). The resulting CSHMM and cell assignments are presented in Fig. 5. As can be seen, similar to the lung data, for this data the assignment of cells to paths generally agrees with their known function. For example, the 0-1-2-6-8 set of paths lead from the embryonic MEF cells (day 0) to d2_intermediate, then d5_intermediate and finally to Neuron cells (day 22). In contrast, paths 0-1-3, while following the initial set of cells up to day 2, leads to a different outcome by day 5 (the d5_failedReprog fate). Other trajectories are likely representing the fact that cells are unsynchronized. For example, the 0-1-5-7 paths represent a slightly less mature set of cells along a reasonable trajectory (embryonic - d2_intermediate - d2_induced-d5_earlyN). Once again, most prior methods for the representation and analysis of time series scRNA-Seq data are unable to accurately represent this branching process (Fig. 3 (a)(b)(c)(d)). Monocle 2 while doing a good job at identifying the major branching between failed and neuron cells, fails to separate the d5_earlyN and d5_failedReprog which are assigned to different paths in our model. Similarly, PCA and GPLVM do not clearly identify the trajectories and tend to mix the successful and unsuccessful differentiated cells. TSNE was also unable to clearly identify the trajectory from d2 and d5 to neurons. We also ran a discrete version of the CSHMM algorithm (Fig. S2). Similar to the results for the lung developmental data, the discrete model largely agrees with the continuous one in terms of the overall topology. However, they differ in some of the cell assignments (for example, the discrete model assigns some of the later d2_induced cells to path 0) and, as we show below it is also less able to identify cell type specific genes.

**Fig. 3.**
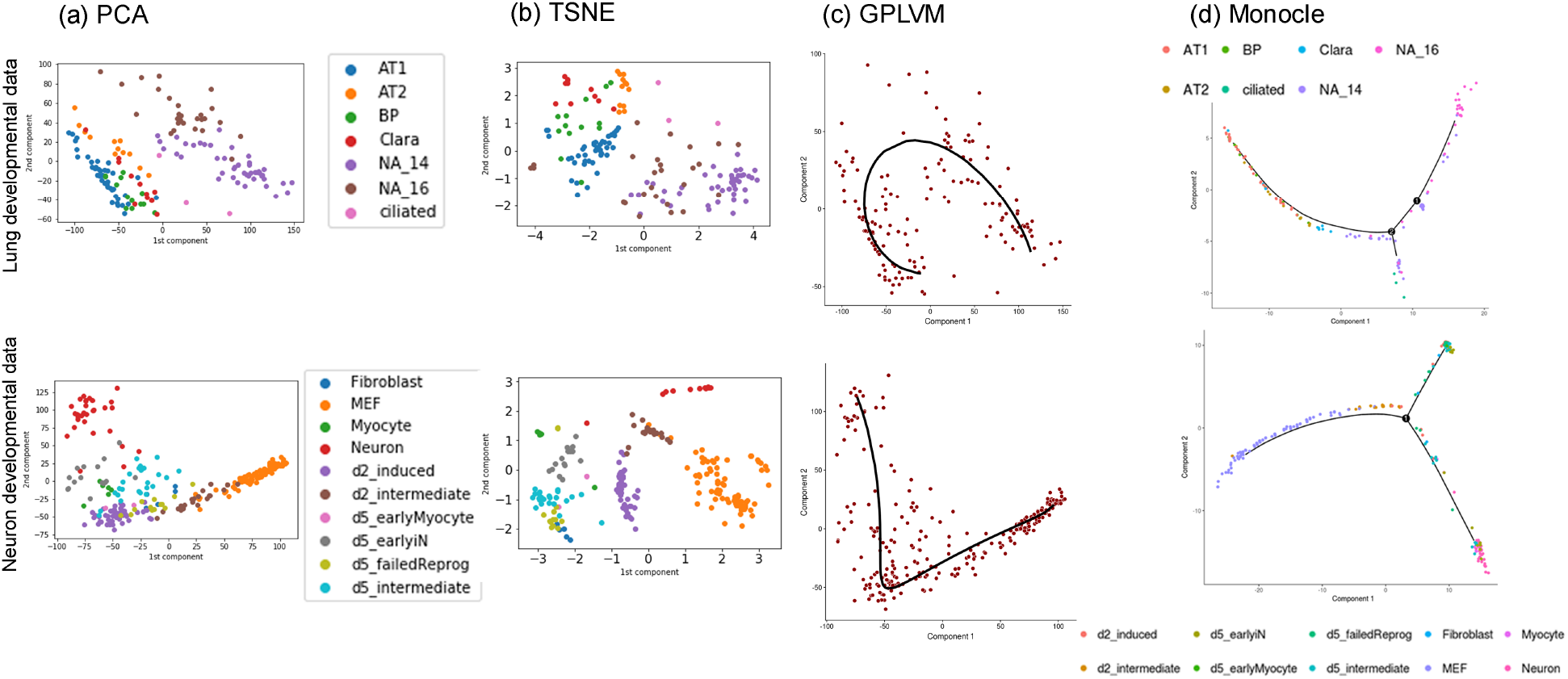
Analysis of lung development and MEF reprogramming data by prior methods. (a) PCA (b) TSNE (c) GPLVM (d) Monocle 2. Top row presents results for the lung dataset and the bottom for the neural developmental dataset. Colors correspond to cell fate assignments in the original papers.

#### 3.2.1 Identifying genes activated during neural cell development

Several known and novel genes can be identified using the continuous cell assignments (Fig. 4 (a)(b)(c)). For example, Insm1 and Myt1l were identified in the original paper as known neuron markers and their CSHMM reconstructed trajectories agree with such roles. Several other genes not identified in the original paper appear to be highly correlated with successful differentiation. For example, Ngfrap1 (Bex3, Fig. 4(b)) has been identified previously as contributing to nerve growth (Calvo et al., 2015). Similarly, prior studies have shown that Mtmr7 is highly expressed in the brain (Mochizuki and Majerus, 2003). Other genes identified highlight the difference between the discrete and continuous models (Fig. 4(c) and Supporting Results). We also analyzed the top GO categories for the set of genes associated with specific fates. Enriched GO categories for genes correlated with each fate agrees well with known functions. For example, for the neuron path (path 8) the top categories are “neuron part” (p-value 1e-29) and “synapse” (p-value 1e-22). For the path that includes neural progenitors (earlyN, path 7) we see an enrichment for “nervous system development” (p-value 5e-14) as well as for several categories and TFs related to cell proliferation (including E2F with a p-value of 3e-22). In sharp contrast, the “failed reprogramming” path (Path 4) is not enriched for any neural activity and is instead enriched for various extracellular matrix categories (p-value 1e-20). See supporting website for the complete list of enriched categories.

**Fig. 4.**
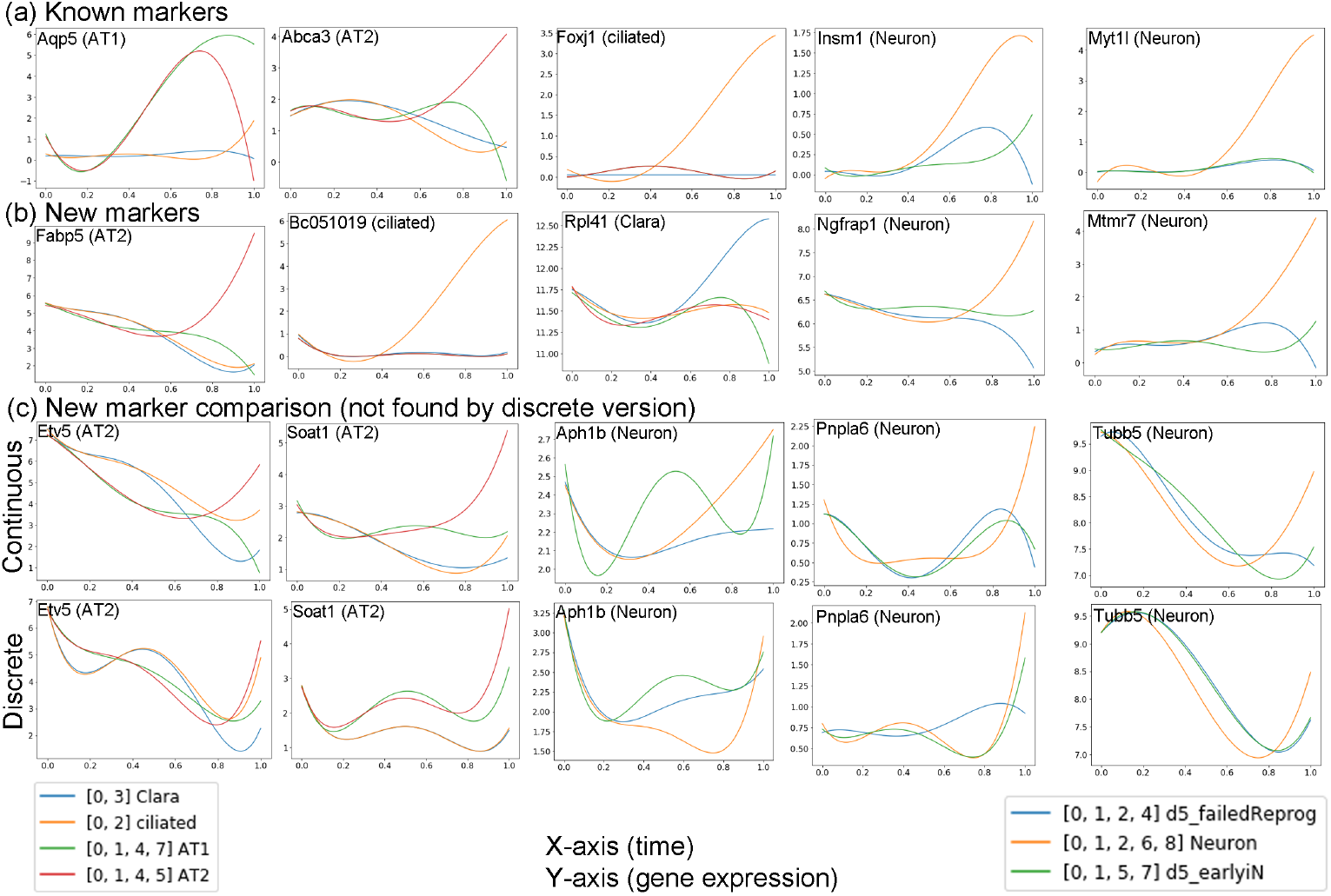
Reconstructed gene expression profiles for lung and neural development data. Each figure plots the expression profile of a gene along the different paths in the corresponding model. Each image includes the gene name and the cell type it was assigned to by the model (AT1, AT2, cillated and Clara from the lung model and Neuron from the neuron model). (Top row) Known markers for the different cell types. (Second row) Novel markers not identified in the original papers found by the CSHMM assignments. (Third and fourth rows) Comparison of reconstructed profiles using the CSHMM (top) and discrete HMM (bottom). Several genes has a unique path profile using the CSHMM but did not display such profile when using the discrete model.

**Fig. 5.**
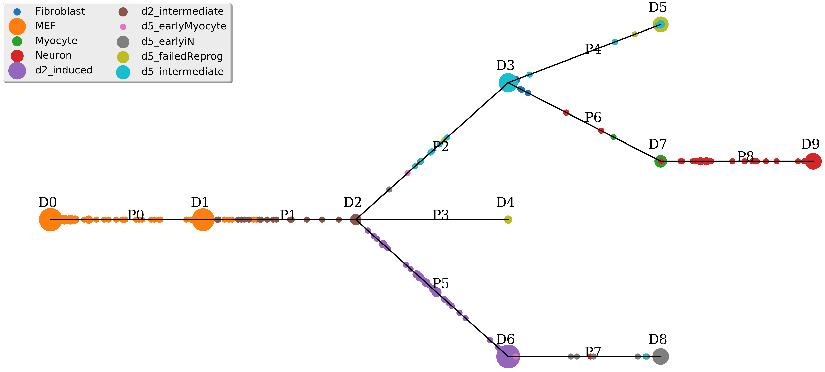
The CSHMM model structure and continuous cell assignment for the MEF reprogramming dataset. Notations, symbols and colors are similar to the ones discussed for Fig. 1.

### 3.3 Scalability and robustness of the CSHMM model

The datasets discussed so far include hundreds of cells with relatively high coverage. Some of the recent scRNA-Seq studies profile a higher number of cells (thousands) though usually with lower coverage. To test the ability of the method to scale to larger number of cells we used both simulated and additional data. For simulated data we re-run the analysis discussed above by replicating each of the 152 lung cells 100 times and adding 20% random dropout to the genes in each replicate (genetating a total of 15K expression profiles). For the CSHMM learning and inference we used the top 1000 most variable genes and tested two versions of search for cell assignment and *K*, either using 10 or 100 values (which is what we used for the smaller dataset). For the 10 values variant, running on a desktop with 4 threads takes roughly 40 minutes to perform one iteration of the EM algorithm and since we usually require less than 10 iterations the total run time is still less than 7 hours even for this dataset. For the 100 values variant the total run time is 15h. The resulting models are presented in Fig. S3 (10 values) and Fig. S4 (100 values). As can be seen, even when using more than 15000 cells, both models reconstruct all the major paths that were recovered in the original (152 cells) lung model. In addition to testing the impact of the number of cells, and different values for the hyper-parameter representing the time points in each path, we also used the simulated data to test the impact of different choices for another hyperparameter of our model, λ_*g*_ which controls the L1 penalty used to select genes for the different paths. As we show (Supplementary section 1.4), we do not observe a large impact on the resulting model for a set of reasonable values for this parameter.

In addition to using simulated data we also tested the CHMM on the 40K cells zebrafish dataset mentioned above. Unlike the two datastes discussed above much less is known about the specific differentiation pathways for several of the last time point cell types. CSHMM analysis of this dataset required only 2 iterations and took 33 hours per iteration. Results are presented in Fig. S24. To determine the success of the assignments focused on the leafs in the model (corresponding to the annotated cell types in the original paper). We calculated the adjusted random index (ARI) agreement between these two sets. We found that the ARI achieved by the CSHMM assignments is significantly better when compared to 1000 randomization tests for the cells (*p* < 10^−10^ based on randomization tests, Fig. S23). Thus, the CSHMM method can scale to larger datasets with tens of thousands of cells.

Another issue that can impact the analysis of scRNA-Seq data is dropout. Due to the lower quantity of RNA obtained from single cells, and the amplification steps required, several genes with low transcript numbers may appear to have 0 transcripts in scRNA-Seq data (Kharchenko et al., 2014). As we discuss in Supporting Results, we performed extensive analysis of dropout impact on the CSHMM method. We observe that for rates which increase dropout by 5-20% results stay largely the same. Beyond 20% additional dropouts we observe a larger impact in which Clara and AT2 cells are merged in a single path though AT1 and AT2 cells are still separated and the initial parts of the model are also correct even for 40% additional dropouts.

## 4 Discussion and future work

Both major strategies for modeling developmental trajectories for time series scRNA-Seq data have advantages and disadvantages. Pseudotime Ordering allows for continuous assignment of cells and the reconstruction of complete expression trajectories. However, these methods often do not take noise into account and the ordering is based on a very limited set of values for each cell. In contrast, probabilistic methods handle noise and the complete set of genes well, but do not provide a continuous representation for the expression profiles.

Here we show that Continuous State HMMs (CSHMMs) can provide a solution for both problems. On the one hand it is a probabilistic method and so can accommodate noise and missing values while on the other hand it provides continuous assignment of cells to paths. We formally defined the CSHMM and discussed methods for learning and inference in such model. We applied our methods to simulated and real scRNA-Seq data. Analysis of the models constructed by the CSHMM method shows that it can accurately reconstruct the branching model for these differentiation processes, correctly assigns cells to the different paths and fates and reconstructs expression trajectories that identified known and novel marker for the different cell types.

While it is impossible to say if the continuous cell assignments orderings determined by the our model are correct (since we do not know the ground truth), a possible way to evaluate the accuracy of these assignments is to look at the resulting gene trajectories. Given a specific ordering, by any method, we can plot the resulting expression profiles for genes in these cells. This can be used to both, identify genes that are in agreement with a specific path in the model and to compare the ordering with orderings obtained by other methods. As we have shown in Fig. 4, genes identified by the CSHMM ordering include several of the known markers for specific cell types, improving upon prior methods. This results provides some support to the accuracy of the cell assignment to paths.

The ability of the CSHMM method to handle noise would be even more important for more recent studies that profile a much larger set of cells with less coverage (and so, with higher noise). As we have shown, CSHMMs can scale to model such data. While these initial results are encouraging, in the future we would like to further improve the efficiency of the learning algorithm so that it can scale to using more genes even with higher cell numbers (for example, by performing sampling on the set of cells used in each EM iteration). We would also like to test the ability to incorporate other types of data, including regulatory information, to aid in improving the model learning and cell assignment.

Software implementing the CSHMM method as well as a README file and an example input are available from the supporting website (http://www.andrew.cmu.edu/user/chiehl1/CSHMM/).

## Footnotes

1 which performs minimum-spanning-tree analysis on a reduced dimension of the data

